# From mass spectral features to molecules in molecular networks: a novel workflow for untargeted metabolomics

**DOI:** 10.1101/2021.12.21.473622

**Authors:** Damien Olivier-Jimenez, Zakaria Bouchouireb, Simon Ollivier, Julia Mocquard, Pierre-Marie Allard, Guillaume Bernadat, Marylène Chollet-Krugler, David Rondeau, Joël Boustie, Justin J.J. van der Hooft, Jean-Luc Wolfender

## Abstract

In the context of untargeted metabolomics, molecular networking is a popular and efficient tool which organizes and simplifies mass spectrometry fragmentation data (LC-MS/MS), by clustering ions based on a cosine similarity score. However, the nature of the ion species is rarely taken into account, causing redundancy as a single compound may be present in different forms throughout the network. Taking advantage of the presence of such redundant ions, we developed a new method named MolNotator. Using the different ion species produced by a molecule during ionization (adducts, dimers, trimers, in-source fragments), a predicted molecule node (or neutral node) is created by triangulation, and ultimately computing the associated molecule’s calculated mass. These neutral nodes provide researchers with several advantages. Firstly, each molecule is then represented in its ionization context, connected to all produced ions and indirectly to some coeluted compounds, thereby also highlighting unexpected widely present adduct species. Secondly, the predicted neutrals serve as anchors to merge the complementary positive and negative ionization modes into a single network. Lastly, the dereplication is improved by the use of all available ions connected to the neutral nodes, and the computed molecular masses can be used for exact mass dereplication. MolNotator is available as a Python library and was validated using the lichen database spectra acquired on an Orbitrap, computing neutral molecules for >90% of the 156 molecules in the dataset. By focusing on actual molecules instead of ions, *MolNotator* greatly facilitates the selection of molecules of interest.

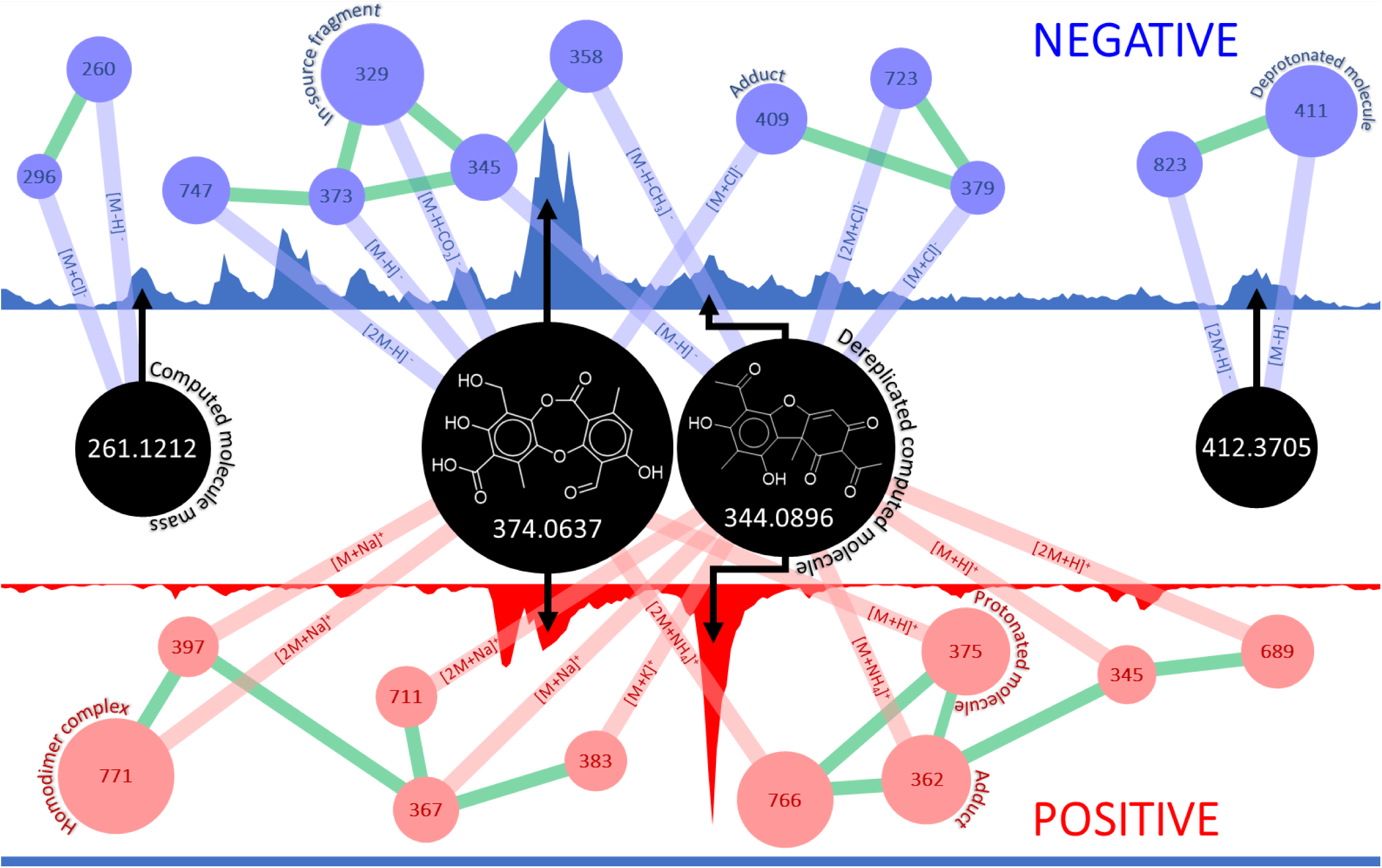

## Introduction

Molecular networks have become a cornerstone of metabolomics analysis approaches, notably since the establishment of the GNPS with the creation of large community-based MS2 databases,^1^ and the Feature-Based Molecular Networking (FBMN) approach, which combines cosine clustering with quantitative comparative metabolomics.^2,3^ In the form of mass spectral networks, the spectral data becomes easier to interpret, with ions being represented by nodes, connected by edges if their spectral similarity calculated by a cosine score exceeds a set threshold. The potential identity of the nodes can be established by dereplication, using the MS2 spectral libraries available on the GNPS, giving insight into the identity of neighbouring nodes using the network topology. The main caveat of this reliance on dereplication is that only a low percentage of the total nodes can be annotated using MS2 databases,^4^ leaving the user with a large amount of unknown nodes. Improvements have been proposed to increase the annotation coverage, including dereplication using in-silico spectral libraries,^5,6^ annotation with shared spectral patterns with MS2LDA,^7^ or with chemical compound classifications enabled by CANOPUS^8,9^ or MolNetEnhancer.^10^ However, most nodes in networks remain without precise dereplication, which can be attributed to different reasons: (i) nodes being originated from truly unknown compounds, (ii) nodes from known compounds absent from databases, (iii) nodes that remain undereplicated due to low cosine scores owing to different acquisition parameters and (iv) nodes that represent known or unknown molecules but from ion species absent from databases. Since the first two points can mostly be solved by a global community effort and that the third one implies standardization across methods and instruments, we here focused on the last point. While in a typical LC-MS based metabolites profiling analysis, a metabolite can produce mass spectral molecular features in the form of various adducts in both positive and negative ionization modes, most spectral libraries contain a majority of [M+H]^+^ ions, which could explain the low annotation rate in networks. Moreover, ion species of a same molecule do not necessarily have similar MS2 spectra, meaning that multiple nodes from a same molecule will form separate clusters. The existence of adduct and fragment nodes in a network implies that the user must often perform manual inspection to locate the redundant ions. To overcome this hurdle and assist in the annotation process, we here introduce a Python library named *MolNotator*, centred on highlighting molecules within the ions constituting mass spectral networks. After running *MolNotator*, molecules are represented as nodes, displayed amidst the ions they produced, in their ionization context (adducts, complexes, fragments, etc.). Within this context, all ions produced by a molecule can be easily spotted, and moreover, by using the neutral nodes as anchors, positive and negative ionization data can be merged into a single network. In its core, *MolNotator* relies on combinatorial triangulations to compute the molecular nodes, allowing the most relevant interpretation of the data given the present ions. Furthermore, using the metadata *MolNotator* computes, a dereplication module was added, with optional retention time and ion species filters to reduce false positive annotations. In addition to MS^2^ dereplication with MGF files, the module can also dereplicate neutral nodes using molecular masses or formulae from user-provided CSV or TSV files. *MolNotator* was validated using a set of 156 purified secondary metabolites from lichen origin that were analysed in Data-Dependent Acquisition (DDA) and manually processed to extract each MS/MS spectra. Using *MolNotator*, 92.3% of the targeted molecules were correctly predicted with an average < 2 ppm mass error. Methods have been developed to deal with adduct annotation^11–15^ and a method similar to this one was recently published, called Ion Identity Molecular Networking (IIMN),^16^ which uses chromatographic shape correlations for clustering instead of combinatorial triangulations. Indeed, the authors of IIMN also report that ion species diversity in databases is lacking and that many nodes in molecular networks are redundant. *MolNotator* improves dereplication possibilities by propagating annotations to the neutral nodes and helps producing databases from scratch with its MGF Updater module: the inputted MGF can be updated with the adduct annotation of each ion and the molecule they belong to. Notably, this method merges positive and negative ionization modes which are typically studied separately. During our validation, most molecules were found in both positive and negative modes, however, running *MolNotator* on experimental data (complex extracts with molecules present at lower abundance) will show how each mode will lead to the detection of their unique molecules. We further demonstrate that dereplication is also improved through the use of *MolNotator* by applying its adduct and retention time filters and by the use of molecular mass for formula-based dereplication. *MolNotator* is available to the community as a Python package.

## Materials and Methods

### Sample preparation

156 molecules from our previous work to produce the Lichen Database were also used here.^17^ These were dissolved in UPLC-grade methanol at a concentration of 10 μg/mL prior to analysis. The molecule list is available in the Supporting Information as a TSV file.

### LC-MS analysis

All molecules were analyzed with a generic data dependent LC-HRMS/MS method in both positive and negative ionization modes. Chromatographic separation was performed on an Acquity UPLC system (Waters, Milford, MA, USA) interfaced to a Q-Exactive Focus mass spectrometer (Thermo Scientific, Bremen, Germany), using a heated electrospray ionisation source (HESI-II). The LC conditions were as follows: column: Waters BEH C18 100 × 2.1 mm, 1.7 μm; mobile phase: (A) Milli-Q water + 0.1% formic acid; (B) UPLC grade acetonitrile with 0.1% formic acid; gradient: 5% B (0 min), 100% B (7 min), 100% B (10 min), 5% B (10.01 min), 5% B (11 min); flow rate: 600 μL/min; injection volume: 2 μL. The optimised parameters of the HESI-II source were: source voltage: 3.5 kV, gas flow rate (N_2_): 48 units; auxiliary gas flow rate: 11 units; reserve gas flow rate: 2.0; capillary temperature: 256.2 °C (pos), RF level of S-Lens: 45. The mass spectrometer was calibrated using a mixture of caffeine, methionine-arginine-phenylalanine-alanine-acetate (MRFA), sodium dodecyl sulphate, sodium taurocholate and Ultramark 1621 in an acetonitrile/methanol/water solution containing 1% formic acid by direct infusion. The data-dependent MS/MS acquisitions were performed on the 3 most intense ions detected in the full scan MS (Top3 experiment). The width of the MS/MS isolation window was 1 Da, and the normalized collision energy (NCE) was set at 15, 30 and 45 units. In the data-dependent MS/MS experiments, full scans were acquired at a resolution of 35,000 FWHM (at m/z 200) and MS/MS scans at 17,500 FWHM with an automatic maximum injection time. After being acquired in the MS/MS scans, the precursor ions were placed in a dynamic exclusion list for 2.0 seconds.

### MZmine 2 parameters

Thermo’s proprietary *raw* files were converted to *mzXML*^18^ format using the MSConvert module^19^ from ProteoWizard.^20,21^ All files were imported in MZmine 2.39^22^ for Linux and were processed using the GenOuest computer cluster located in Rennes, France, using an Intel(R) Xeon(R) Gold 5220 CPU, 2.20GHz. The parameters used are available in the Supporting Information (**Table S1**). The ADAP chromatogram builder and deconvolution modules were used.^23^ The processed ion data was exported in MGF format and the attributes for each feature in CSV format.

### MolNotator Python package, dependencies, and parameters used

The MGF and CSV files obtained from MZmine were placed in the mzmine_out folder the *MolNotator* project before starting the process. To use *MolNotator*, it is important to note that each sample file (“sample.mzXML” for example) should be prefixed “POS_” and “NEG_”, respectively for the positive and negative ionization data, and aside from the prefix, the sample should have the same name in both modes (*i*.*e*. “POS_sample.mzXML” and “NEG_sample.mzXML”), so that they can be linked during the mode merging process. The main libraries *MolNotator* relies on are matchms 0.6.2,^24^ pandas 1.3.5^25^ and numpy 1.19.4.^26^ The networks are produced using Cytoscape 3.7.1^27^ by importing the node and edge tables produced by *MolNotator*. All parameters are described and available in the parameter file provided with the dataset on Zenodo (https://doi.org/10.5281/zenodo.5792605). They are also transcribed here *pro forma*. Duplicate filter: mass error = 0.002 Da, retention time error = 10 s. Fragnotator: matched peaks = 3, score threshold = 0.1, mass error = 0.001 Da, retention time error = 10 s. Adnotator: mass error = 0.002 Da, precursor mass error = 0.1 Da, retention time error = 10 s, cosine threshold = 0.2, hard cosine threshold = 0.6, adducts searched= 21 in POS, 22 in NEG (see SI) run BNR (Basic Neutral Requirements, see SI) = True, BNR NEG = [“[M-H]-”, “[2M-H]-”], BNR POS = [“[M+H]+”, “[M+Na]+”, “[M+NH4]+”, “[2M+H]+”, “[2M+Na]+”, “[2M+NH4]+”]. Mode merger: mass error = 0.003 Da, precursor mass error = 0.1, retention time error = 15 s, BNR NEG = [“[M-H]-”, “[2M-H]-”], BNR POS = [“[M+H]+”, “[M+Na]+”, “[M+NH4]+”, “[2M+H]+”, “[2M+Na]+”, “[2M+NH4]+”]. MGF updater: skipped. Dereplicator: ion dereplication was carried out using the LDB^17^ (cosine threshold = 0.5, matched peaks = 2, precursor error = 0.004 Da, mass error = 0.002 Da, retention time error = 10 s, top hits = 3, adduct filter = True, retention time filter = True). Neutral dereplication was made using a TSV file produced by merging the COCONUT^28^ and LOTUS^29^ databases (precursor mass error = 0.001 Da, top hits = 3). *MolNet* parameters: mass error = 0.002 Da, cosine threshold = 0.98, matched peaks = 8.

### Data and Code availability

The *MolNotator* package can be downloaded from [GitHub](https://github.com/ZzakB/MolNotator) and [Pypi](https://pypi.org/project/MolNotator/). We suggest creating a new Python environment prior to using *MolNotator*. The package is provided with a small sample of the data for testing purposes, and the full dataset is available on Zenodo (https://doi.org/10.5281/zenodo.5792605). Two project folders are provided, one prior to the use of *MolNotator* and one after processing, containing the final networks. Both come with a style folder for Cytoscape to fit each network type.

## Results and discussion

### Lichen Database Orbitrap spectra extraction and manual curation

Spectra from the files of the 156 analytes were extracted semi-automatically as done in our previous work^17^ and added to the GNPS servers in addition to the other Lichen Database spectra already present there ([negative](https://gnps.ucsd.edu/ProteoSAFe/gnpslibrary.jsp?library=LDB_NEGATIVE) and [positive](https://gnps.ucsd.edu/ProteoSAFe/gnpslibrary.jsp?library=LDB_POSITIVE) ionization modes). Doing so, 335 anion and 367 cation adducts were collected, the molecules which produced them were used to validate *MolNotator*. This dataset will be referred to hereafter as LDB-Orbitrap.

### *MolNotator* glossary of terms

*MolNotator* functions as a pipeline composed of five main modules: *Fragnotator* which detects in-source fragments, *Adnotator* which detects adducts produced by a single molecule and calculates the accurate mass of that molecule, *Mode merger* which merges negative and positive mode data based on the predicted molecules, their accurate masses and their retention time, *Dereplicator* which dereplicates the data with user-provided databases (MGF, CSV or TSV) and *Cosiner* which adds a layer of cosine-based clustering to the network. Cosine calculations done throughout the processing use the matchms library^24^). **Figure 1** presents the elements composing a *MolNotator* cluster.

**Figure 1:**
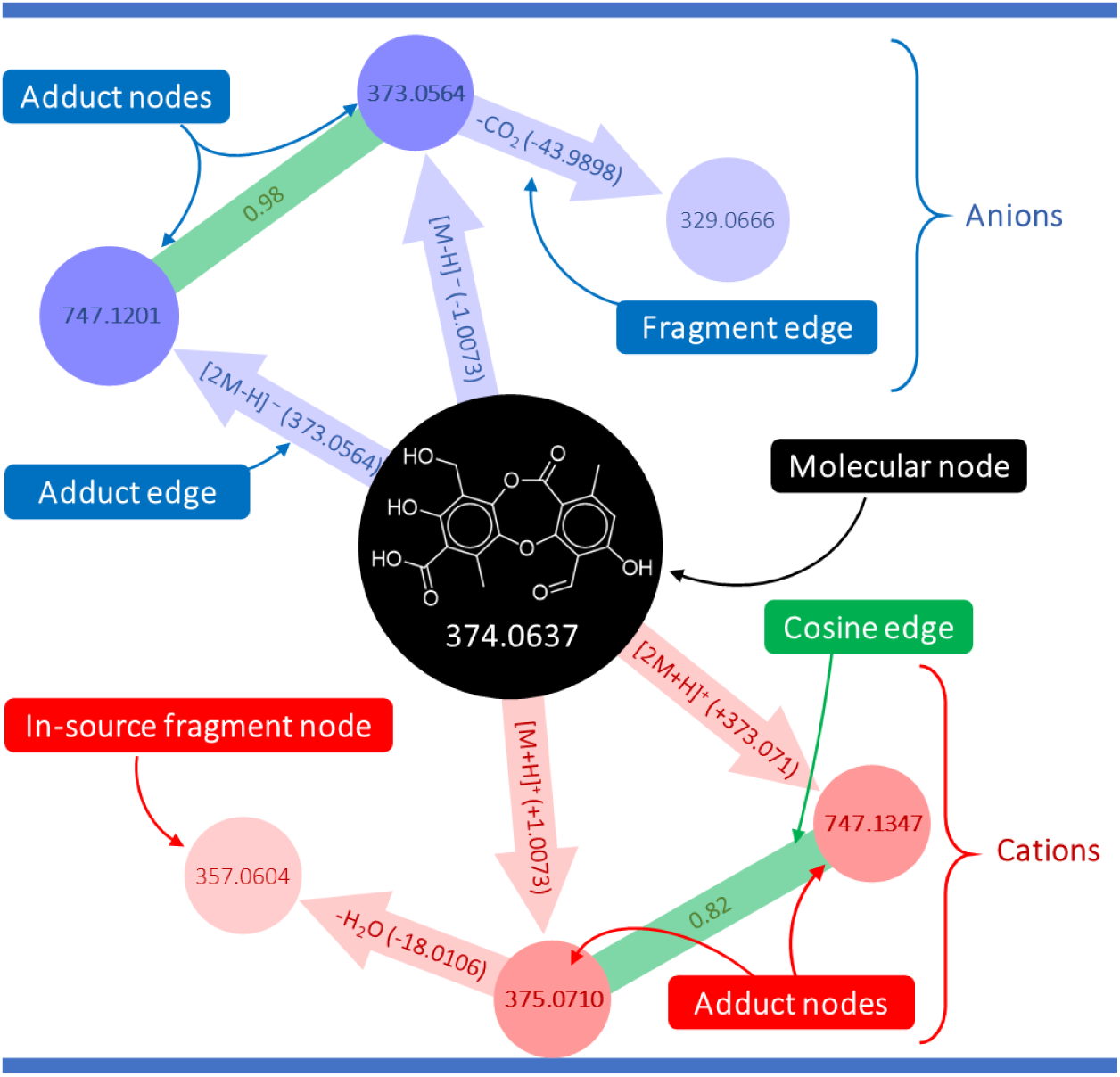
Elements of a *Molnotator* network.

#### Predicted molecule

a molecular mass, computed by combinatorial triangulation using the *m/z* values of ions detected in mass spectrometry. At least two *adducts* are necessary for *Adnotator* to triangulate a molecular mass, otherwise the single ion is not processed. They are represented in the networks by *Molecular nodes*.

#### Molecular node

node representing a *predicted molecule*, connected to all ions used for its calculation. The *predicted molecule* they represent can be dereplicated by propagating the annotations from the nodes directly connected. See the black node in **Figure 1**.

#### Adducts

protonated and deprotonated molecules along with all ion species used to compute the molecular masses. These can be anions or cations, and also dimers, trimers or any formula provided by the user in the *adduct table*. See the blue and red nodes connected to the black node in **Figure 1**.

#### In-source fragment

a molecule’s fragment generated in the ion source that was intense enough to be picked by the DDA and fragmented like any other ion. It is therefore associated to an MS/MS spectrum and appears in regular networks like any other ion. See the pink and light blue nodes in **Figure 1**.

#### Adduct edge

edge connecting an *adduct* node to their *molecular node* (edges connecting the molecular node to the adduct nodes in **Figure 1**). No spectral similarity is calculated for these, as they connect an MS/MS spectrum to a mass.

#### Fragment edge

edge connecting an in-source fragment node to their in-source precursor node.

#### Cosine edge

edge connecting two nodes as long as their cosine similarity score is above threshold (green edges in **Figure 1**).

#### Adduct table

table provided by the user with all adduct formulae to be searched by *MolNotator*.

#### Ionization hypothesis

an ion species formula from the *adduct table* assigned to a given ion. *Ionization hypotheses* are used by *Adnotator* to gauge which adduct that ion is more likely to be and compute the associated molecule.

#### Hypothetical molecule

a calculated molecular mass for an ion given its *m/z* value and its *Ionization hypothesis*. One or more of these *hypothetical molecules* are calculated for each ion by *Adnotator* before selecting the most likely candidate.

#### Basic Neutral Requirements (BNR)

a list of adducts, of which at least one should be detected for a hypothetical molecule to be validated. Molecules displaying none of these ions are discarded.

The modules are further expanded upon in the Supporting Information, illustrated by the clustering of glomellic acid ions. This typical lichen didepside helps underlining the importance of clustering ions based on the ionization context, as it is prone to producing several in-source fragments in addition to its adducts in both ionization modes.

### *MolNotator* performance

Processing all files through *MolNotator* using an Intel(R) Core(TM) i7-6700HQ CPU 2.60GHz on a laptop took 17 hours and 46 minutes without any form of parallel computing, which is not yet implemented (**Figure 2**). As a reference point, processing all the files on MZmine using an Intel(R) Xeon(R) Gold 5220 CPU 2.20GHz on the computer cluster and parallel calculations took about the same time. The two main bottlenecks are *Adnotator*, notably for positive mode files, and *MolNet*. The run time for *Adnotator* increases with the number of files processed and more importantly with the number of ions in each file, especially if there are many adduct species to search like in this experiment. *MolNet* high run times are dependent on the total number of ions as the module calculates cosine scores between each ion pair for each neutral.

**Figure 2:**
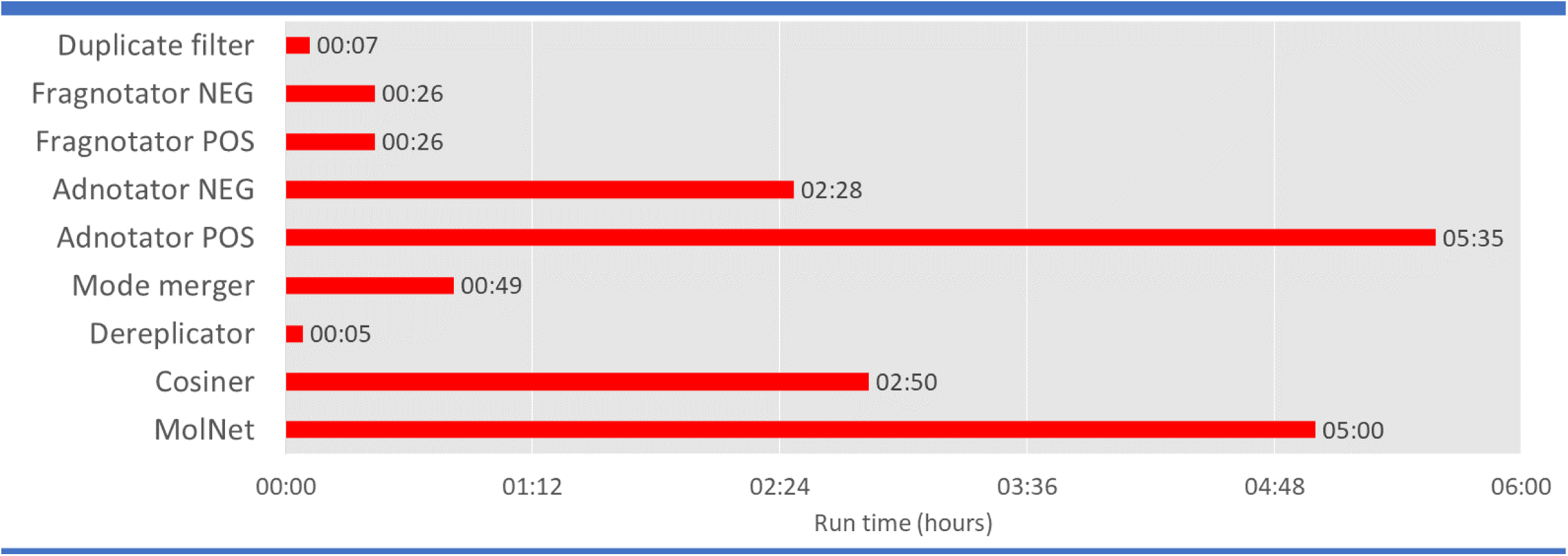
Run time for *MolNotator* for each module in hours.

### *MolNotator* simplified networks

*MolNotator* produces three types of networks, named *MolNetworks* 1 (all ions and neutrals), 2 (adducts and neutrals only) and 3 (neutrals only clustered by cosine similarity). See **Figures 3 and 4**. Overall, the clustering resulting of *MolNotator* simplifies molecular networks, as demonstrated in **Figure 3**, representing networks generated using the glomellic acid MGF file, one of the 156 LDB analytes. After each *MolNotator* step, the user can chose to export sample-wise networks in addition to the network for all processed samples. In this case, the glomellic acid positive and negative ionization mode networks produced by cosine similarity are compared to the “mixed” *MolNetworks* 1 and 3. All nodes belonging to the metabolite are contained within a single cluster (circled in purple) in the *MolNetwork* 1. In the negative and positive mode networks, the same nodes (colored yellow and circled purple) are spread throughout several clusters and can even be found as singletons. Moreover, in such networks, typically, hydrogen, sodium and potassium adducts from a same molecule will not be contained in the same cluster using spectrum-based similarities, as they often have different fragmentation patterns. In addition to that, in-source fragments may form separate clusters, something that is common with depside-type molecules which are split in the mass spectrometer ion source due to the fragile ester bond. The *MolNetwork* 1 is comparatively smaller: 575 nodes compared to the 642 of the two other regular networks (260 and 382 in negative and positive ionization modes respectively), here a 10% size reduction (mixed network on the left in **Figure 3**). Furthermore, considering the *MolNetwork* 3 (**Figure 3**, lower right) only 175 neutral nodes remain out of the 575 nodes from the *MolNetwork* 1. This represents a 72% network size reduction.

**Figure 3:**
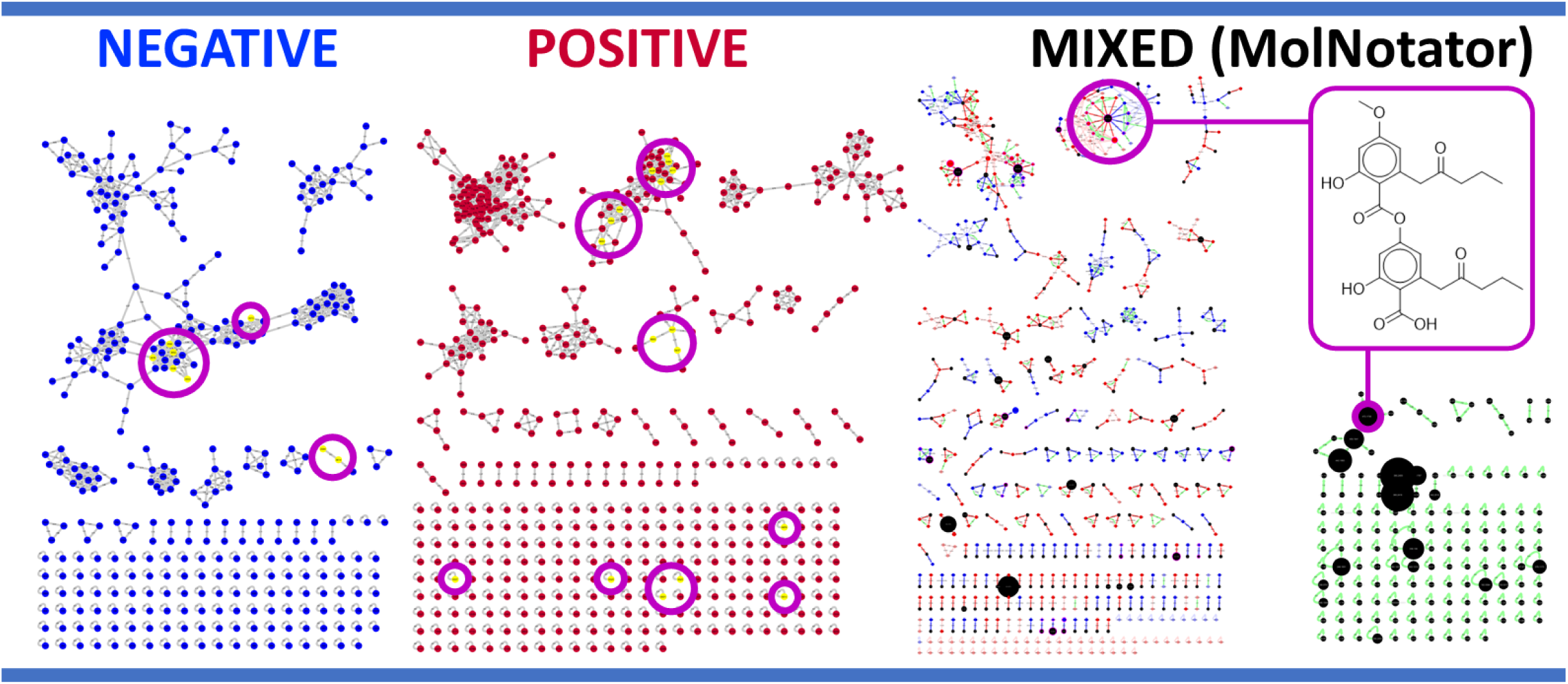
Comparison between cosine-based molecular networks (negative and positive modes) and the *MolNotator* networks (mixed, *MolNetwork* 1 with all ions and neutrals on the left, and on the right the *MolNetwork* 3 displaying only neutral nodes connected by cosine edges) for the glomellic acid files. Nodes belonging to glomellic acid are colored yellow and circled purple in the first two networks. These nodes are all contained in the purple circle in the *MolNetworks*. Many other molecules were detected in the glomellic acid file, mostly contaminants and closely related lichen compounds like glomelliferic acid.

**Figure 4:**
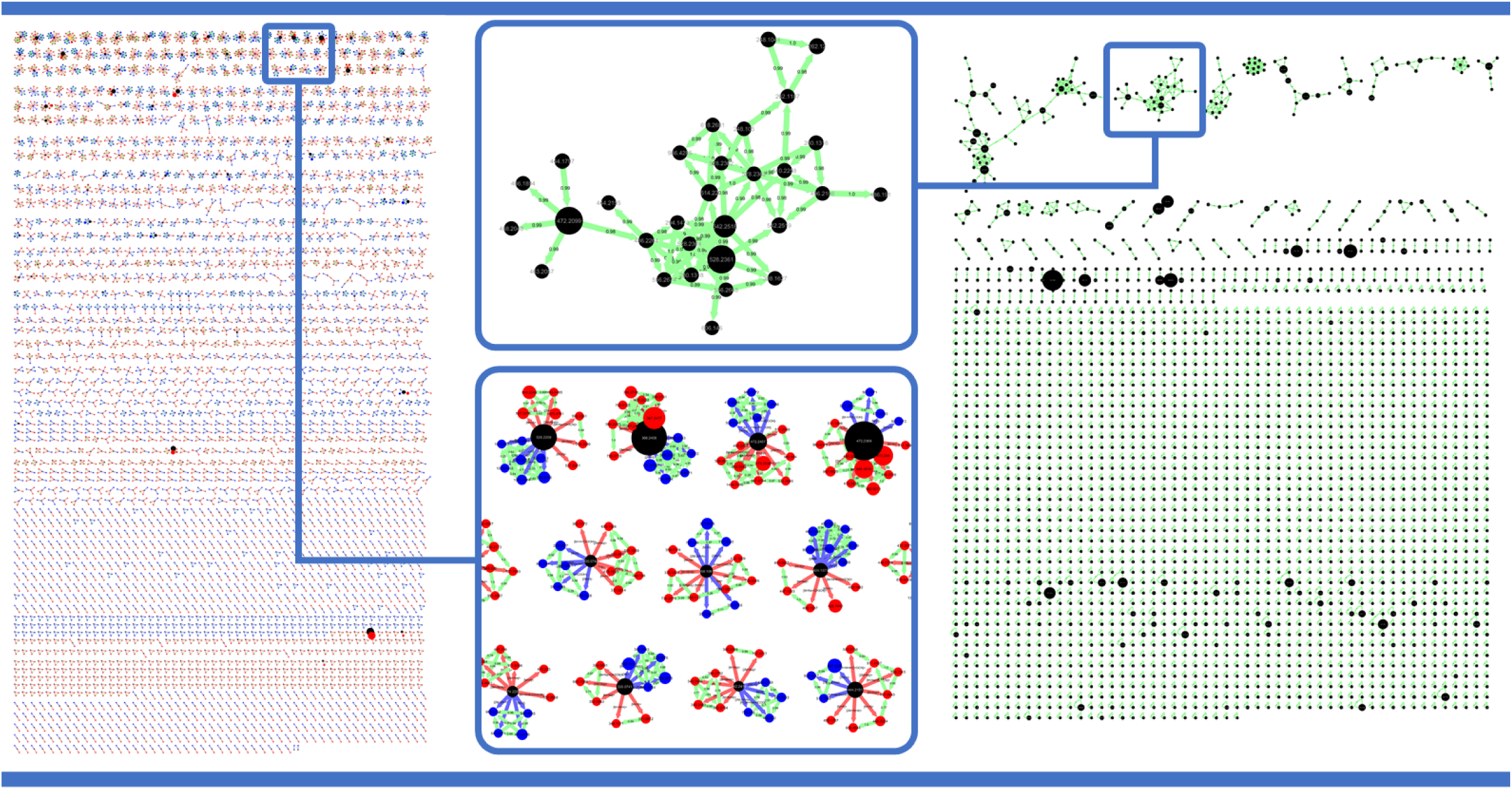
Network produced by the *MolNet* module. The network on the left represents the 2544 neutral nodes along with their adducts directly connected to them. The one on the right represents the same 2544 neutrals without the adducts but connected through cosine edges. Most of these molecules are unrelated to the 156 analytes, being mostly derivatives, degradation products and contaminants.

Zooming out from one specific molecule and considering all the obtained mass spectrometry data from all the files processed, the *MolNetworks* 2 (only molecular and adduct nodes) and 3 (neutrals only) were produced and displayed in **Figure 4**. By ignoring in-source fragment, precursor and singleton nodes, the first network shows what is in essence a list of all predicted molecules, whether they are the 156 searched metabolites or any other molecule predicted in the data. Most metabolites from the LDB are present in the top of the network, amongst the nodes with the highest adduct count. The *MolNetwork* 3 removes all ions and focuses on cosine similarities between neutral nodes (based on their omitted adducts). The low degree of clustering observed here is caused by the high thresholds used for *MolNet* in order to avoid too large clusters (cosine threshold = 0.98, matched peaks = 8). All networks produced can be found on Zenodo (https://doi.org/10.5281/zenodo.5792605).

### Method validation

Out of the 156 metabolites, 144 (92.3%) were correctly represented as molecular nodes. The average mass error, highly dependent on the instrument and its calibration, was 1.8 ppm, with 70% of compounds being calculated within a 0 to 2 ppm range (**Figure 5A**). A dereplication was carried out using LDB-Orbitrap files now available on the GNPS servers. 137 out of the 144 predicted molecules were thus dereplicated. As *MolNotator* can also be used to extract spectra and produce databases, and by comparison to the manually curated LDB-Orbitrap library, the method retrieved 529 anion adducts and 649 cation adducts, compared to the corresponding 335 and 367 already in the library.

**Figure 5:**
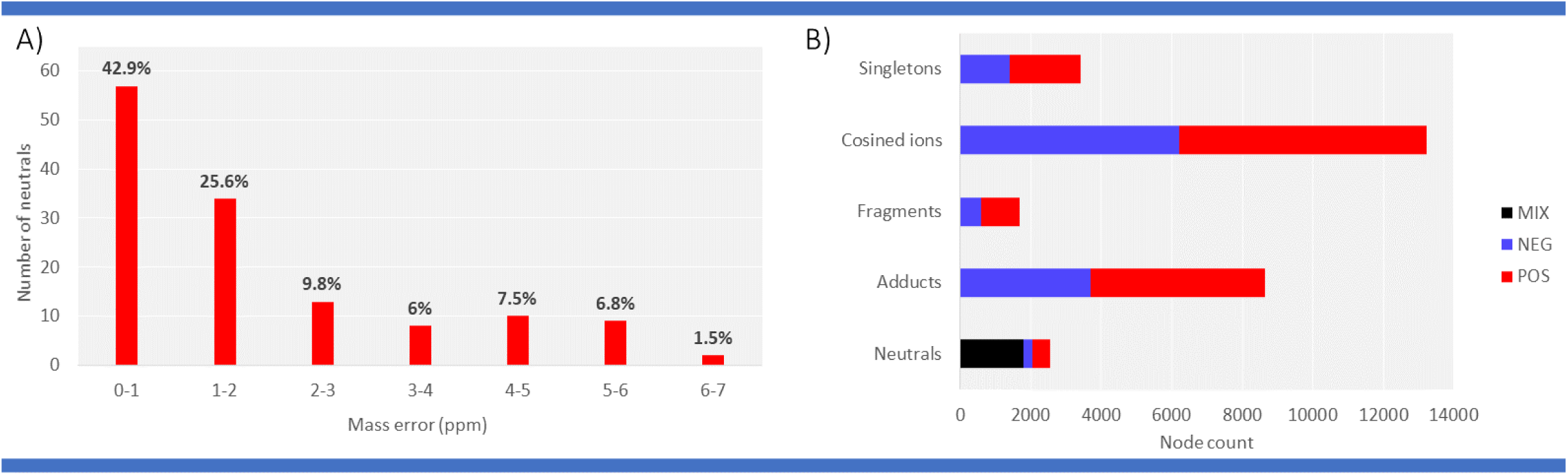
A: Mass precision in ppm of the 144 *MolNotator* predicted molecules in the mass spectral data. The neutral node mass is calculated by averaging the hypothetical molecule mass of each ion connected to that neutral. B: adduct abundance in the network generated (considering the 156 analytes and all the other molecules detected). The “POS” and “NEG” sections of the Neutral bar represent neutral nodes found specifically in positive and negative ion modes respectively.

### Adduct abundance patterns

The global network contained 2544 molecular nodes, 26 957 ions including 8637 adducts and 1693 in-source fragments. The number of molecules computed by *MolNotator* is largely superior to the number of analytes expected (156). Most of these can be considered to be contaminants, derivatives or degradation products from the analytes, showcasing how much can be detected even when analysing “pure” standards. When considering all 2544 molecules, each produced on average 4 ions (adducts or in-source fragments, **Figure 5B**). Most of the other ions were clustered using cosine scores, leaving only 3404 singleton nodes (12.6% of ions). When considering only the 144 metabolites from the LDB that were correctly computed, the ion per molecule ratio is of 8, which can be attributed to the fact that those were highly concentrated and thus highly abundant, thereby producing more (visible) ions. The ion species abundance was calculated considering all 2544 molecules and the 144 analytes separately, and they both follow the same trends (**Figure 6A and B**).

**Figure 6:**
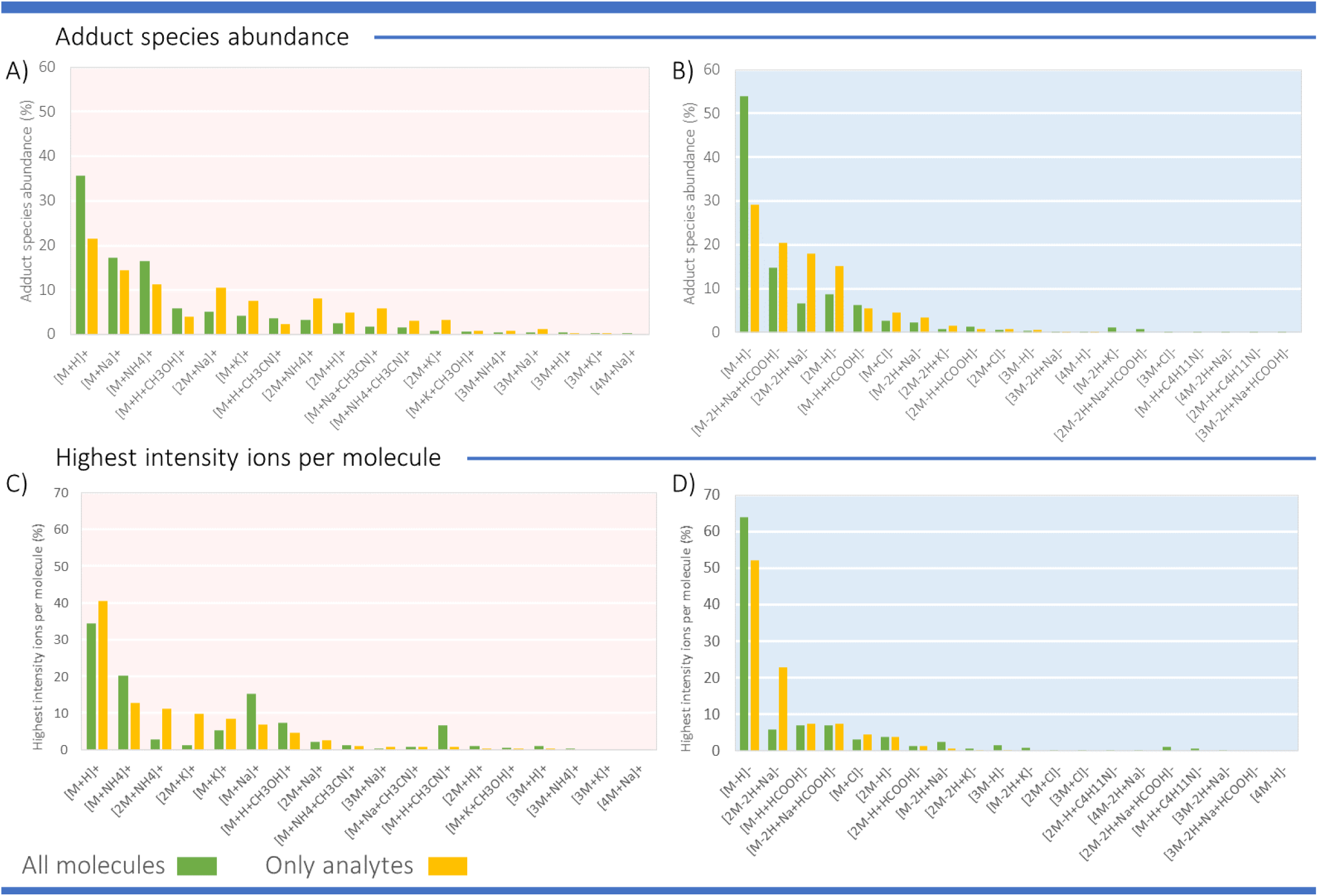
A and B: adduct species abundance in positive (pink background) and negative (lightblue background) ionization modes respectively. C and D: most highly ionized species per molecule in positive and negative ionization modes respectively. “All molecules” refers to all the 2544 compounds computed (analytes and contaminants) and “Only analytes” refers to the 144 found metabolites from the LDB.

Considering only the 144 analytes, for the positive ionization mode (**Figure 6A**), [M+H]^+^ were indeed the most abundant species, accounting for 21.5% of all adducts formed, followed by [M+Na]^+^ (14.5%), [M+NH_4_]^+^ (11.2%) and [2M+Na]^+^ (10.4%). In negative ionization mode (**Figure 6B**), [M-H]^-^ species also were the most produced species (29.1%) followed by [M-2H+Na+HCOOH]^-^ (20.4%), [2M-2H+Na]^-^ (17.9%) and [2M-H]^-^ (15.2%). Although (de)protonated species are indeed the most abundant, most ions are still to be found in so called degenerate features, which are quite often overlooked.^30^ These other species account for 78.5 and 70.9% of ions here, respectively in positive and negative ionization modes. It is remarkable that adducts like [M-2H+Na+HCOOH]^-^ are seldom considered in adduct annotation software, since it is the second most abundant ion in negative ionization mode in this experiment, yet virtually absent from spectral libraries. It could be argued that these degenerate features, although often detected, represent only minor ions close to the analytical noise, while (de)protonated molecules are the major visible peaks in most LC-MS chromatograms. However, this statement could also be challenged: in negative ionization mode (**Figure 6D**), [M-H]^-^ was the most intense ion produced in 64% of cases, while in positive ionization mode (**Figure 6C**) the protonated molecule was the most intense species in only 34% of occurrences, in competition with [M+NH_4_]^+^ and [M+Na]^+^ which were the most abundant ionized species in 20% and 15% of occurrences, respectively (**Figure 6C and D**). Thus, it is fair to state that (de)protonated molecules do not represent the overwhelming majority of ions encountered in LC-MS data, nor are they necessarily the most abundant ionized species for each molecule. Moreover, not taking into account the ionization context of molecules is a major contributing factor to the low dereplication rates in molecular networks as these adducts are absent from spectral databases. These results emphasize the importance of clustering redundant features from molecules, something *MolNotator* can do as part of metabolomics analysis workflows.

## Discussion

*MolNotator* represents LC-MS/MS data as true molecular networks, with predicted neutrals linked to all adducts and in-source fragments that are associated to them, whether they are formed in negative or positive ionization mode. The method was validated using spectra from the LDB, 92.3% of which were correctly predicted, thereby retrieving 1178 spectra compared to the 702 already collected manually, due to the annotation of additional ion species for these LDB molecules. In addition to providing context for the ions in a mass spectral network, *MolNotator* has proven to be an efficient way of producing a spectral database from raw data, as the additional spectra can be simply pulled from the original data (MGF file) and added to the library, along with their dereplication and adduct annotation. However, 12 molecules could not be correctly predicted as neutral nodes, mostly due too a low adduct count (0 or 1 ion produced, see Supporting information). While the data tested here was from pure standards, where the target analyte is highly concentrated, *MolNotator* can also function with complex samples. The library already contains functionality that can triangulate compounds between files: in a sample in which a molecule would be represented by a single ion, thus making triangulation of the neutral impossible, it will extrapolate the knowledge from other samples where more ions were produced to compute the neutral. Hence, in the network of the first sample, that single ion will be connected nonetheless to its molecular node. This makes the method highly effective when processing large metabolomics datasets, in which the number of samples will help the overall triangulations.

Regarding the run time, *MolNotator’s* main bottleneck is *Adnotator*, which was capable of processing samples containing 300 ions in 1 minute, and was tested in other experiments with samples containing several thousand spectra and processing them in a dozen minutes. Future improvements should include the use of parallel computing and graph theory to greatly enhance performance on *Adnotator* and *MolNet* as well.

Besides making insightful networks, our analysis also highlights the amount of redundant data in LC-MS/MS experiments and the implications it has on dereplication. This point was also stressed in the IIMN publication for Ion Identity Networks, also suggesting that databases should integrate more adduct species.^16^ The main differences of *MolNotator* compared to IIMN are the calculation methods used (combinatorial triangulations instead of chromatographic shape correlations), the use of molecular nodes in the networks which include accurate mass dereplications, and the merging of positive and negative ionization modes that resulted in networks combining both ionization modes. The use of precursor-based triangulations can prevail if the chromatographic peaks are too low quality because of the low(er) abudance of the metabolite and/or the low MS^1^ to MS/MS ratio typically observed during DDA analyses. Thus, *MolNotator* offers a complementary approach that is independent of chromatographic shapes. The use of the neutral nodes for dereplication make it easier to create MS/MS libraries when most information available is in the form of compound (structure) databases, which is common with understudied organisms, like lichens. The use of both complementary ionization modes in the same network allows for a more comprehensive assessment of the sample’s chemical content, instead of focusing on only one ionization mode. As such, *MolNotator* facilitates the more widespread use of dual ionization acquisition polarities and the combined data processing, analysis and interpretation.

The use of the ionization context in mass spectral networks allows researchers to focus on the molecules themselves, providing the user with a better rationale to understand the nodes displayed. Furthermore, it makes the implementation of an adduct filter possible, thereby improving the quality of dereplication using spectral libraries. The various ions connected to the neutral nodes can serve as replicate spectra for a single molecule, and may help tools like CANOPUS and MolNetEnhancer determining its chemical compound class.^8,10^

Given its good scalability, in future projects, we plan to apply *MolNotator* to complex metabolite extracts to annotate as many molecular nodes as possible using the literature, the biochemical insights and ion species context provided by the *MolNotator* network. We further aim to map the ion species distributions in (public) experimental data akin to the distributions displayed of the validation dataset. We will also create additional ion species-enriched spectral libraries to gain a more comprehensive overview of the impact of ion species on metabolomics data. Finally, using the *MolNotator* ion species assignments, we would like to investigate the mass spectral network creation based on a unique ion species type, such as a protonated or deprotonated species. This would yield further insights in the true extent of unique chemicals metabolomics data can capture and how they relate to each other.

We conclude that by using *MolNotator*, metabolomics researchers can directly deal with molecules instead of mass spectral features or ions, and thus get a glimpse of the samples’ the full molecular content. *MolNotator* can easily be combined with other metabolomics preprocessing, processing, and analysis tools to form an integral part of untargeted metabolomics workflows aiming to decomplexify mass spectral networks.

## Supporting information

Molecule list

## Acknowledgement

We acknowledge the GenOuest bioinformatics core facility (https://www.genouest.org) for providing the computing infrastructure. We thank Rennes Métropole for the “Mobilité sortante” grant to support the collaboration with J. L. Wolfender’s laboratory. We also thank R. Lücking and H. Sipman (Botanic Garden and Botanical Museum in Berlin) for giving us access to samples from S. Huneck’ chemical library and J. A. Elix (Canberra) for standards from his own collection.

## Supporting Information Available

- Molecule_list.tsv: list of all molecules used in this experiment.
- Zenodo dataset (https://doi.org/10.5281/zenodo.5792605): raw and processed data used.

## Notes

https://doi.org/10.5281/zenodo.5792605

### Competing Interest Statement

The authors have declared no competing interest.

