## Supporting Information for "From mass spectral features to molecules in molecular networks: a novel workflow for untargeted metabolomics"

‖*Univ Paris-Saclay, CNRS, BioCIS UMR 8076, F-92290 Châtenay-Malabry, France*

⊥*Univ Rennes, IETR UMR 6164, F-35000, Rennes, France*

#*Univ Bretagne Occidentale, Département de Chimie, F-29238, Brest, France*

@*Bioinformatics Group, Wageningen University, Wageningen, the Netherlands*

### MZmine parameters

Table S1: MZmine parameters. When different, positive ionization mode (POS) and negative ionization mode (NEG) parameters are both stated.

| **Module** | **Parameters** |
|---|---|
| Raw data methods<br>>Mass detection | Scans: MS level: 1<br>Mass detector: centroid<br>Noise level: 5.0E5 (POS), 2.0E5 (NEG)<br>Mass list name: masses |
| Raw data methods<br>>Mass detection | Scans: MS level: 2<br>Mass detector: centroid<br>Noise level: 0<br>Mass list name: masses |
| Peak list methods<br>>Peak detection<br>>ADAP Chromatogram builder | Scans: MS level: 1<br>Mass list: masses<br>Min group size in # of scans: 5<br>Group intensity threshold: 5.0E5 (POS), 2.0E5 (NEG)<br>Min highest intensity: 5.0E5 (POS), 2.0E5 (NEG)<br>m/z tolerance: 15 ppm |

**Table S1 continued from previous page**

| Module | Parameters |
|---|---|
| Peak list methods<br>>Peak detection<br>>Chromatogram deconvolution | Algorithm: Wavelets (ADAP):<br>S/N threshold: 10<br>S/N estimator: Wavelet Coeff. SN:<br>Peak width mult.: 3<br>abs(wavelet coeffs.): checked<br>min feature height: 5.0E5 (POS), 2.0E5 (NEG)<br>coefficient/area threshold: 150<br>Peak duration range: 0.00-1.00<br>RT wavelet range: 0.00-0.20<br>m/z center calculation: MEDIAN<br>m/z range for MS2 scan pairing (Da): 0.03<br>RT range for MS2 scan pairing (min): 0.1 |
| Peak list methods<br>>Isotopes<br>>Isotopic peaks grouper | m/z tolerance: 8.0ppm<br>Retention time tolerance: 0.08 absolute (min)<br>Monotonic shape: unchecked<br>Maximum charge: 2<br>Representative isotope: Lower m/z |
| Peak list methods<br>>Alignment<br>>Join aligner | m/z tolerance: 15.0ppm<br>Weight for m/z: 1<br>Retention time tolerance: 0.2 absolute (min)<br>Weight for RT: 1<br>Require same charge state: unchecked<br>Require same ID: unchecked<br>Compare isotope pattern: unchecked |

**Table S1 continued from previous page**

| Module | Parameters |
|---|---|
| Peak list methods | Keep only peaks with MS2 scan (GNPS): checked |
| >Filtering | Reset the peak number ID: checked |
| >Peak list rows filter | (rest is unchecked / unused) |
| | Mass list: masses |
| | Merge MS/MS (experimental): checked |
| | Select spectra to merge: across samples |
| | m/z merge mode: weighted average (remove outliers) |
| | intensity merge mode: sum intensities |
| Peak list methods | Expected mass deviation: 5.0ppm |
| >Export/Import | Cosine threshold (%): 70.0 |
| >Export for/Submit to GNPS | Peak count threshold (%): 40.0 |
| | Isolation window offset (m/z): 0.0 |
| | Isolation windows width (m/z): 3.0 |
| | Filter rows: ALL |
| | Submit to GNPS: unchecked |
| | Open folder: Unchecked |

**Table S1 continued from previous page**

| Module | Parameters |
|---|---|
| Peak list methods<br>>Export/Import<br>>Export to CSV file | Field separator: “,”<br>Export common elements:<br>Export row ID: checked<br>Export row m/z: checked<br>Export row retention time: checked<br>Export data file elements:<br>Peak area: checked<br>Export quantitation results: unchecked<br>Identification separator: “;”<br>Filter rows: ALL |

### ***MolNotator*** adduct tables.

Table S2: Positive mode primary adduct table. M: "M" count in the formula, C: Complexity, G: adduct group.

| Adduct_code | Adduct | Charge | Mass | M | C | G |
|---|---|---|---|---|---|---|
| M1\|p1H\|pCH3CN | [M+H+CH3CN]+ | 1 | 42.034374 | 1 | 3 | H |
| M1\|p1H\|pCH3OH | [M+H+CH3OH]+ | 1 | 33.03404 | 1 | 3 | H |
| M1\|p1H\| | [M+H]+ | 1 | 1.007825 | 1 | 0.5 | H |
| M1\|p1NH4\|pCH3CN | [M+NH4+CH3CN]+ | 1 | 59.060923 | 1 | 3 | H |
| M1\|p1NH4\| | [M+NH4]+ | 1 | 18.034374 | 1 | 2 | H |
| M1\|p1Na\|pCH3CN | [M+Na+CH3CN]+ | 1 | 64.016319 | 1 | 4 | Na |
| M1\|p1Na\| | [M+Na]+ | 1 | 22.98977 | 1 | 2 | Na |
| M1\|p1K\|pCH3OH | [M+K+CH3OH]+ | 1 | 70.989923 | 1 | 3 | K |
| M1\|p1K\| | [M+K]+ | 1 | 38.963708 | 1 | 3 | K |

**Table S2 continued from previous page**

| Adduct_code | Adduct | Charge | Mass | M | C | G |
|---|---|---|---|---|---|---|
| M2\|p1H\| | [2M+H]+ | 1 | 1.007825 | 2 | 3 | H |
| M2\|p1NH4\| | [2M+NH4]+ | 1 | 18.034374 | 2 | 3 | H |
| M2\|p1Na\| | [2M+Na]+ | 1 | 22.98977 | 2 | 4 | Na |
| M2\|p1K\| | [2M+K]+ | 1 | 38.963708 | 2 | 4 | K |
| M3\|p1H\| | [3M+H]+ | 1 | 1.007825 | 3 | 4 | H |
| M3\|p1NH4\| | [3M+NH4]+ | 1 | 18.034374 | 3 | 4 | H |
| M3\|p1Na\| | [3M+Na]+ | 1 | 22.98977 | 3 | 5 | Na |

Table S3: Positive mode secondary adduct table. M: "M" count in the formula, C: Complexity, G: adduct group.

| Adduct_code | Adduct | Charge | Mass | M | C | G |
|---|---|---|---|---|---|---|
| M3\|p1K\| | [3M+K]+ | 1 | 38.963708 | 3 | 5 | K |
| M4\|p1H\| | [4M+H]+ | 1 | 1.007825 | 4 | 5 | H |
| M4\|p1NH4\| | [4M+NH4]+ | 1 | 18.034374 | 4 | 5 | H |
| M4\|p1Na\| | [4M+Na]+ | 1 | 22.98977 | 4 | 6 | Na |
| M4\|p1K\| | [4M+K]+ | 1 | 38.963708 | 4 | 6 | K |

Table S4: Negative mode primary adduct table. M: "M" count in the formula, C: Complexity, G: adduct group.

| Adduct_code | Adduct | Charge | Mass | M | C | G |
|---|---|---|---|---|---|---|
| M1\|m1H\|pC4H11N | [M-H+C4H11N]- | -1 | 72.081324 | 1 | 3 | H |
| M1\|m1H\|pHCOOH | [M-H+HCOOH]- | -1 | 44.997655 | 1 | 3 | H |
| M1\|m1H\| | [M-H]- | -1 | -1.007825 | 1 | 1 | H |
| M1\|p1Cl\| | [M+Cl]- | -1 | 34.968853 | 1 | 2 | Cl |
| M1\|m2Hp1Na\|pHCOOH | [M-2H+Na+HCOOH]- | -1 | 66.9796 | 1 | 5 | H |

**Table S4 continued from previous page**

| Adduct_code | Adduct | Charge | Mass | M | C | G |
|---|---|---|---|---|---|---|
| M1\|m2Hp1Na\| | [M-2H+Na]- | -1 | 20.97412 | 1 | 4 | H |
| M1\|m2Hp1K\| | [M-2H+K]- | -1 | 36.948058 | 1 | 4 | H |
| M2\|m1H\|pC4H11N | [2M-H+C4H11N]- | -1 | 72.081324 | 2 | 4 | H |
| M2\|m1H\|pHCOOH | [2M-H+HCOOH]- | -1 | 44.997655 | 2 | 4 | H |
| M2\|m1H\| | [2M-H]- | -1 | -1.007825 | 2 | 3 | H |
| M2\|p1Cl\| | [2M+Cl]- | -1 | 34.968853 | 2 | 3 | Cl |
| M2\|m2Hp1Na\|pHCOOH | [2M-2H+Na+HCOOH]- | -1 | 66.9796 | 2 | 6 | H |
| M2\|m2Hp1Na\| | [2M-2H+Na]- | -1 | 20.97412 | 2 | 5 | H |
| M2\|m2Hp1K\| | [2M-2H+K]- | -1 | 36.948058 | 2 | 5 | H |
| M3\|m1H\| | [3M-H]- | -1 | -1.007825 | 3 | 4 | H |
| M3\|p1Cl\| | [3M+Cl]- | -1 | 34.968853 | 3 | 4 | Cl |

Table S5: Negative mode secondary adduct table. M: "M" count in the formula, C: Complexity, G: adduct group.

| Adduct_code | Adduct | Charge | Mass | M | C | G |
|---|---|---|---|---|---|---|
| M3\|m2Hp1Na\|pHCOOH | [3M-2H+Na+HCOOH]- | -1 | 66.9796 | 3 | 7 | H |
| M3\|m2Hp1Na\| | [3M-2H+Na]- | -1 | 20.97412 | 3 | 6 | H |
| M4\|m1H\| | [4M-H]- | -1 | -1.007825 | 4 | 5 | H |
| M4\|p1Cl\| | [4M+Cl]- | -1 | 34.968853 | 4 | 5 | Cl |
| M4\|m2Hp1Na\|pHCOOH | [4M-2H+Na+HCOOH]- | -1 | 66.9796 | 4 | 8 | H |
| M4\|m2Hp1Na\| | [4M-2H+Na]- | -1 | 20.97412 | 4 | 7 | H |

### Terms used in *MolNotator* networks.

**Predicted molecule**: a molecular mass, computed by combinatorial triangulation using the $m/z$ values of ions detected in mass spectrometry. At least two *adducts* are necessary for *Adnotator* to triangulate a molecular mass, otherwise the single ion is not processed. They are represented in the networks by *Molecular nodes*.

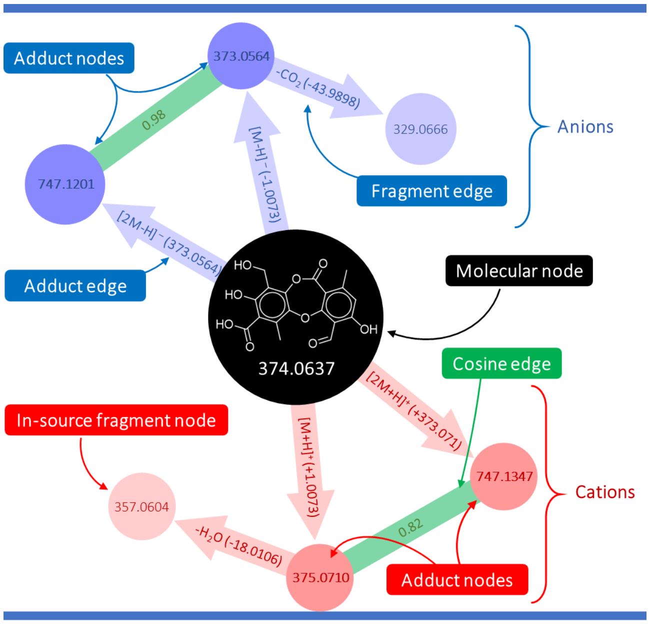


Figure S1: Elements of a *Molnotator* network.

**Molecular node**: node representing a *predicted molecule*, connected to all ions used for its calculation. The *predicted molecule* they represent can be dereplicated by propagating the annotations from the nodes directly connected. See the black node in **Figure S1**.

**Adducts**: protonated and deprotonated molecules along with all ion species used to compute the molecular masses. These can be anions or cations, and also dimers, trimers or any formula provided by the user in the *adduct table*.See the blue and red nodes connected to the black node in **Figure S1**.

**In-source fragment**: a molecule's fragment generated in the ion source that was intense enough to be picked by the DDA and fragmented like any other ion. It is therefore associated to an MS/MS spectrum and appears in regular networks like any other ion. See the pink and light blue nodes in **Figure S1**.

**Adduct edge**: edge connecting an *adduct* node to their *molecular node* (edges connecting the molecular node to the adduct nodes in **Figure S1**). No spectral similarity is calculated for these, as they connect an MS/MS spectrum to a mass.

**Fragment edge**: edge connecting an in-source fragment node to their in-source precursor node.

**Cosine edge**:edge connecting two nodes as long as their cosine similarity score is above threshold (green edges in **Figure S1**).

**Adduct table**: table provided by the user with all adduct formulae to be searched by *MolNotator*.

**Ionization hypothesis**: an ion species formula from the *adduct table* assigned to a given ion. *Ionization hypotheses* are used by *Adnotator* to gauge which adduct that ion is more likely to be and compute the associated molecule.

**Hypothetical molecule**: a calculated molecular mass for an ion given its *m/z* value and its *Ionization hypothesis*. One or more of these *hypothetical molecules* are calculated for each ion by *Adnotator* before selecting the most likely candidate.

**Basic Neutral Requirements (BNR)**: a list of adducts, of which at least one should be detected for a hypothetical molecule to be validated. Molecules displaying none of these ions are discarded.

### *MolNotator* modules.

**Duplicate filter.** Filters out duplicate ions for a given retention time and *m/z* value, when this was not done beforehand.

**Fragnotator.** This module connects in-source fragments - precursor pairs by taking as input an *MGF* file outputted by MZmine. For an ion to be considered an in-source fragment of a given precursor, it must fulfill certain requirements based on the following conditions (with the default values used here):

- $\Delta$RT between ions below threshold (10 seconds).
- Fragment's *m/z* value found in the precursor's MS/MS spectrum (at 0.001 Da).
- Both spectra share a minimum number of peaks ($min\_shared\_peaks = 3$).

- Jaccard similarity coefficient calculated above a given threshold ($matching\_score$ = 0.1).

Figure S2: In-source fragment detection by *Fragnotator*. Here, two in-source fragments are produced from glomellic acid, detected as regular ions by the quadrupole and fragmented by the collision cell to register their MS/MS spectra on the Orbitrap. *Fragnotator* detects them as in-source fragments candidates because their ion's $m/z$ is present in glomellic acid's MS/MS spectrum. Given that they share enough peaks with their in-source precursor, they will be annotated as in-source fragments

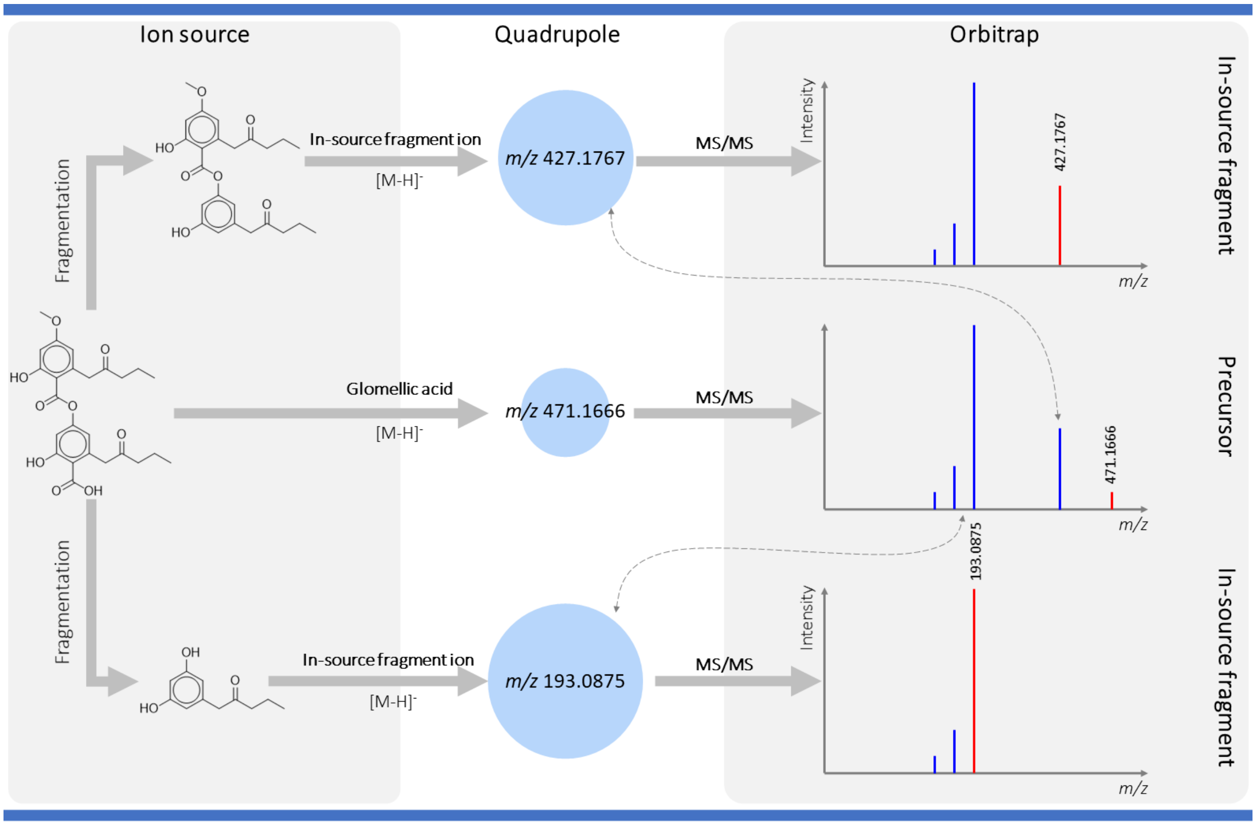


An example is displayed in **Figure S2** to illustrate how *Fragnotator* operates. This module also annotates some edges with common neutral losses, like -$H_2O$, -CO, -$CO_2$, -2$H_2O$, etc. A network example featuring glomellic acid is displayed in **Figure S3**, with the compound's (de)protonated molecules (NEG $m/z$ 471.1663 and POS $m/z$ 473.1814) associated to several in-source fragments or other precursor ions. At this point, it is important to note that these (de)protonated molecules are annotated as in-source fragments: adducts with higher $m/z$ values generally contain these ions in their MS/MS spectrum, making the [M-H]$^-$ and

[M+H]$^+$ virtually indistinguishable from fragments. A good example of how *Fragnotator* rationalises the data can be observed in the positive mode cluster. The protonated molecule's ester bond is fragmented, producing an in-source fragment at $m/z$ 235. That fragment will itself produce its own in-source fragments, either by an $H_2O$ loss ($m/z$ 217) or by a CO loss ($m/z$ 207). All these produce the fragment $m/z$ 189, annotated -HCOOH from $m/z$ 235, but which is more likely to be a succession of -$H_2O$ and -CO as can be observed by the two other fragments.

Figure S3: Clusters produced around (de)protonated molecules of glomellic acid by *Fragnotator*. The [M-H]$^-$ and [M+H]$^+$ ions respectively have $m/z$ values of 471.1663 and 473.1814.

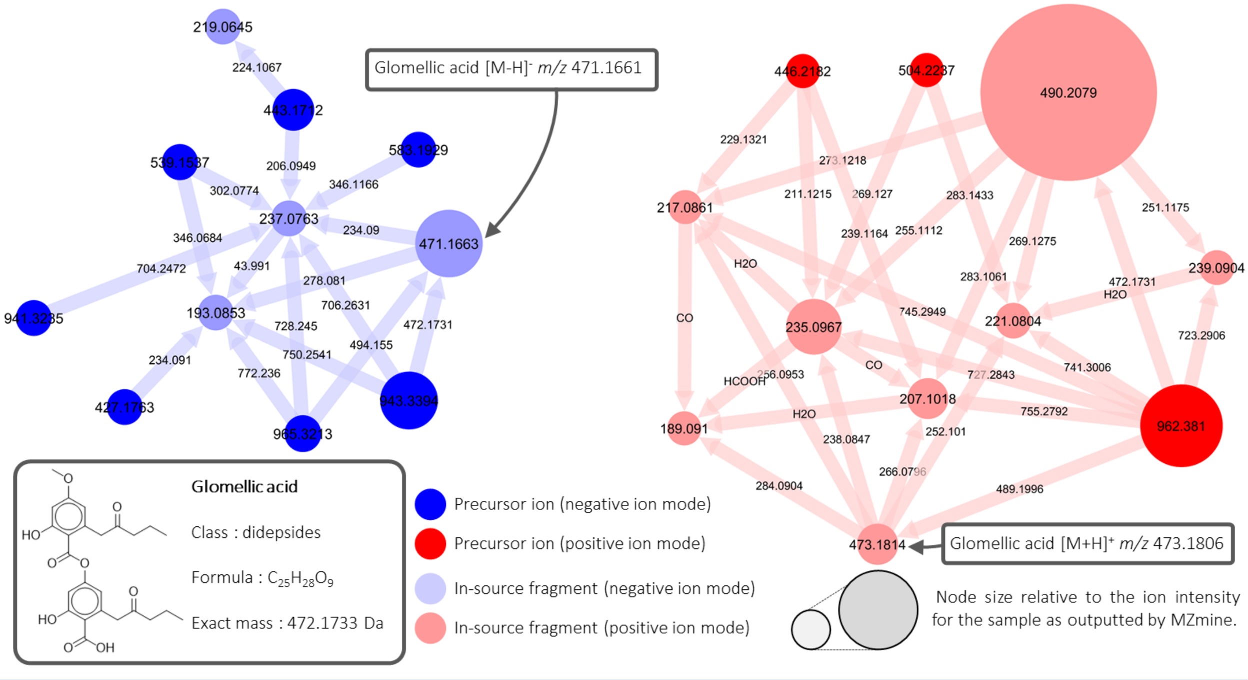

Figure S4: Method used by *Adnotator* to triangulate molecules by their molecular masses and link them up with the adducts they produced.

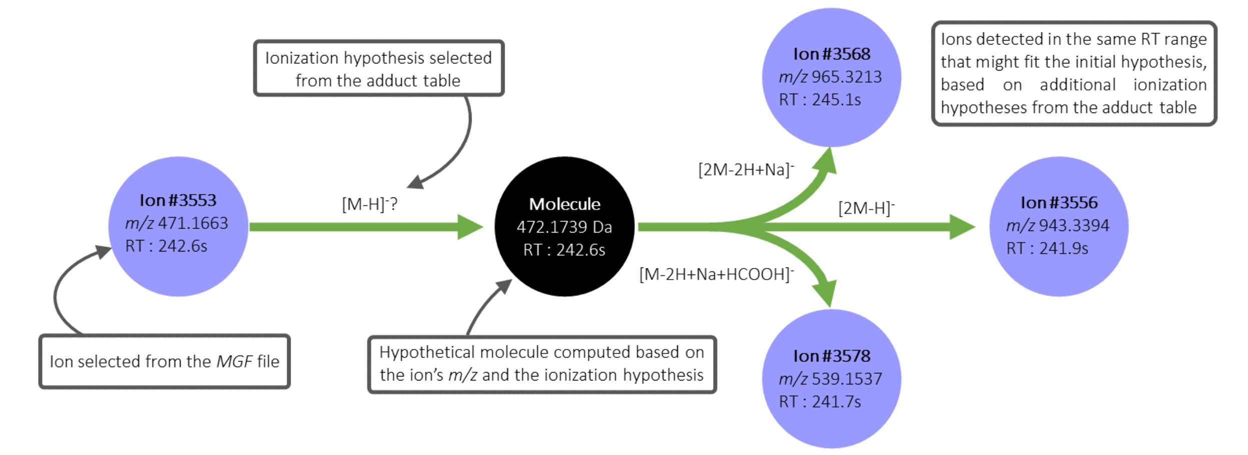


**Adnotator.** This module detects the different adducts produced by a single molecule, representing that molecule as a neutral node linked to all its ions. The basic principle behind this method is that molecules will often produce several ions, in the form of predictable adducts (i.e. $[M+H]^+$, $[2M+H]^+$, $[M+Na]^+$ etc...), the $m/z$ of which can be used to triangulate that molecule's mass. The molecules with the most ions associated are the most likely to exist. The script is provided with two tables of adducts to be searched, as in **Tables S4, S5, S2, S3**. The first table is the primary table, which contains adducts to be used to triangulate the neutral nodes. Adducts in the secondary table are searched in the data once the neutrals are already computed, and they are not used in the triangulations. The primary table should contain all the most common adducts, and the secondary the most rare. This division is made to avoid false positives, reduce computing time and avoid combinatorial explosion. Ions in the *MGF* file are subjected to all ionization hypotheses made available in the primary table. In each case, a hypothetical molecule mass is calculated based on the ion's $m/z$ and the adduct selected. If that molecule exists, it is likely to have produced other ions. In the case displayed in **Figure S4**, if the selected ion is an $[M\text{-}H]^-$, a molecule with a molecular mass of around 472.1739 Da should exist. If that molecule exists, it should have produced other signals, here in the form of $[2M\text{-}H]^-$ $m/z$ 943.3394, $[2M\text{-}2H+Na]^-$ $m/z$ 965.3213 and

[M-2H+Na+HCOOH]$^-$ $m/z$ 539.1537, all detected within the ΔRT window of 10 seconds. Once validated by at least two adducts and cosine scores between adducts exceeding a set threshold, the predicted molecules are represented by nodes with sizes proportional to the summed intensities of all connected adducts. Unlike in regular molecular networks, these molecular nodes do not represent MS/MS spectra. BNR (Basic Neutral Requirements) can be set to filter out unlikely hypothetical molecules. **Figure S5** displays results for glomellic acid after using *Adnotator*. Glomellic acid was correctly computed in both negative and positive ionization modes, respectively at 1.2 and 0.2 ppm.

Figure S5: Clusters produced by *Adnotator* by triangulating molecular masses and associated adducts to their computed molecules.

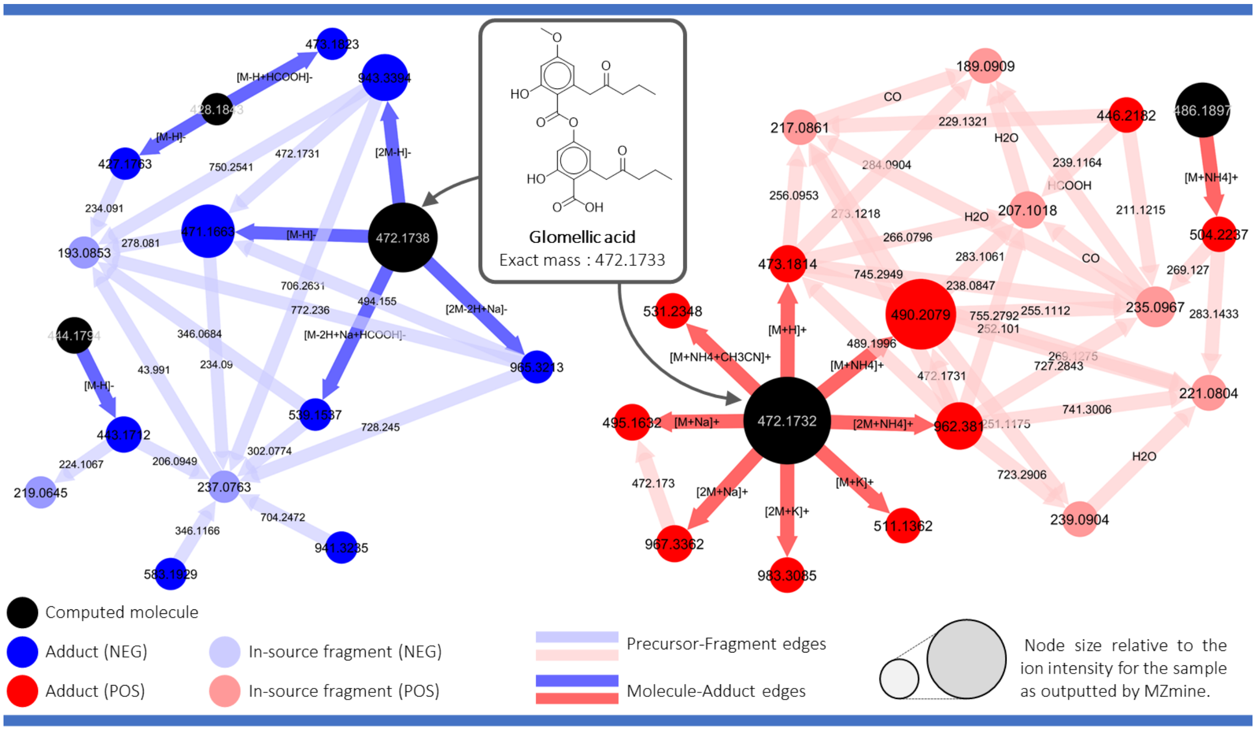


**Mode merger.** This module merges negative and positive ionization mode data based on the computed molecules, under the assumption that two molecules, one in each mode, measured using the same chromatographic conditions, eluting at a similar retention time, with the same calculated mass, are in fact the same molecule. When merged, the annotations from both modes are propagated to the new molecular node. When a molecule is only detected in one ionization mode, a single ion search will be carried out in the opposite mode. Typically, for a molecule displaying $[M+H]^+$ and $[2M+H]^+$ in positive mode, this will search for a single ion in negative mode, generally $[M-H]^-$, which could not produce the molecule by itself because it needs at least another ion to triangulate. Similarly, Mode merger will attempt to generate molecules based on singleton ions from positive and negative modes, producing neutral nodes connected to a single cation and a single anion. These clusterings cannot be cosine validated and to avoid spurious results, BNR can be applied to search only the most likely adducts.

The case of glomellic acid is displayed in **Figure S6**. Instead of having two networks, a single network is produced with all glomellic acid ions in the same cluster. The mass of each molecule is averaged between the neutrals of each mode, having glomellic acid calculated at 0.4 ppm.

Figure S6: Negative and positive ion mode network merging by *Mode merger*.

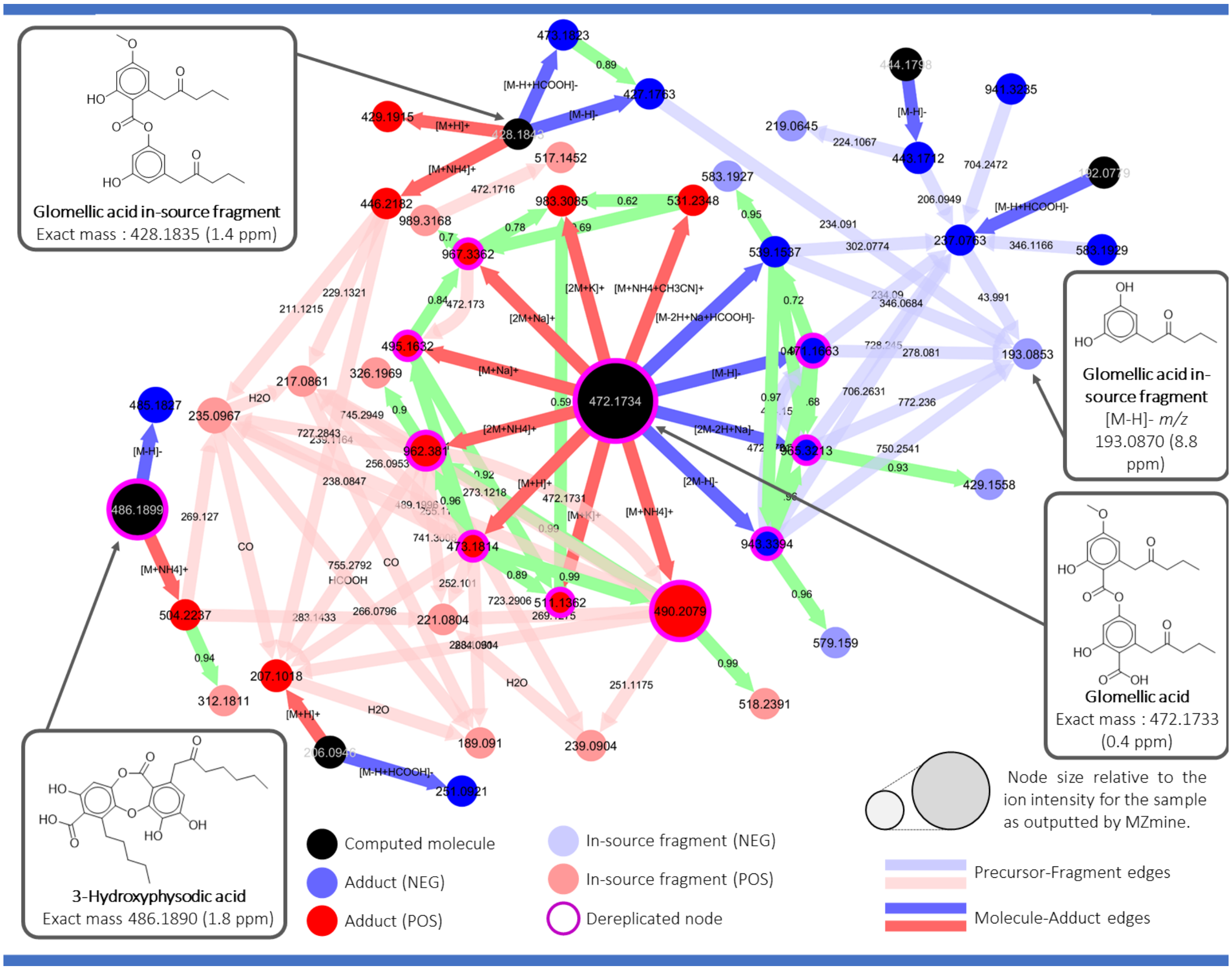


**MGF updater.** Updates the MGF file with the ion species associated with each ion, the molecule they are associated with and the molecular mass and formula calculated by SIRIUS (if available).

**Dereplicator.** This module takes user-provided databases, in MGF, CSV or TSV format, to dereplicate the network data. In the case of MGF files, which contain spectral data, each ion node is compared to the database's spectra and the top hits (as defined by the user) are kept. Among the filters proposed, are the ion mode filter (which is mandatory), a retention time filter and an adduct filter, to keep only hits eluting at the same retention time and displaying the same adduct formula. In the case of TSV or CSV files, molecular nodes are dereplicated using the computed exact mass or formula. Where spectral dereplications may have some shortcomings (molecule absent from the database, low cosine match), the dereplication by TSV/CSV files can compensate. All ion dereplications are propagated to the connected molecular nodes.

**Cosiner.** This module calculates cosine similarity scores between adducts of a same molecule and also between singleton nodes and other nodes, to create cosine edges if the score is above the user-set threshold. This layer of cosine-based clustering is designed to connect ions which could not be associated to their molecular node, to other molecules to which they could be structurally related.To avoid cluttering the network, *Cosiner* does not create edges between neutral nodes. This is left as an option for the last module: *MolNet*.

**MolNet.** Simplifies the network by creating to other mass spectral networks. The first one displays only molecular nodes and adducts, omitting all other nodes to display a list of predicted molecules and their supporting adducts. The second one only displays molecular nodes, connected to each other by cosine edges if two molecules share adducts with high cosine similarity scores.

#### Molecules missed by *MolNotator*.

12 metabolites could not be computed by *MolNotator*, displayed in **Figure S7**.

Figure S7: Molecules that were not predicted as neutral nodes by *MolNotator*

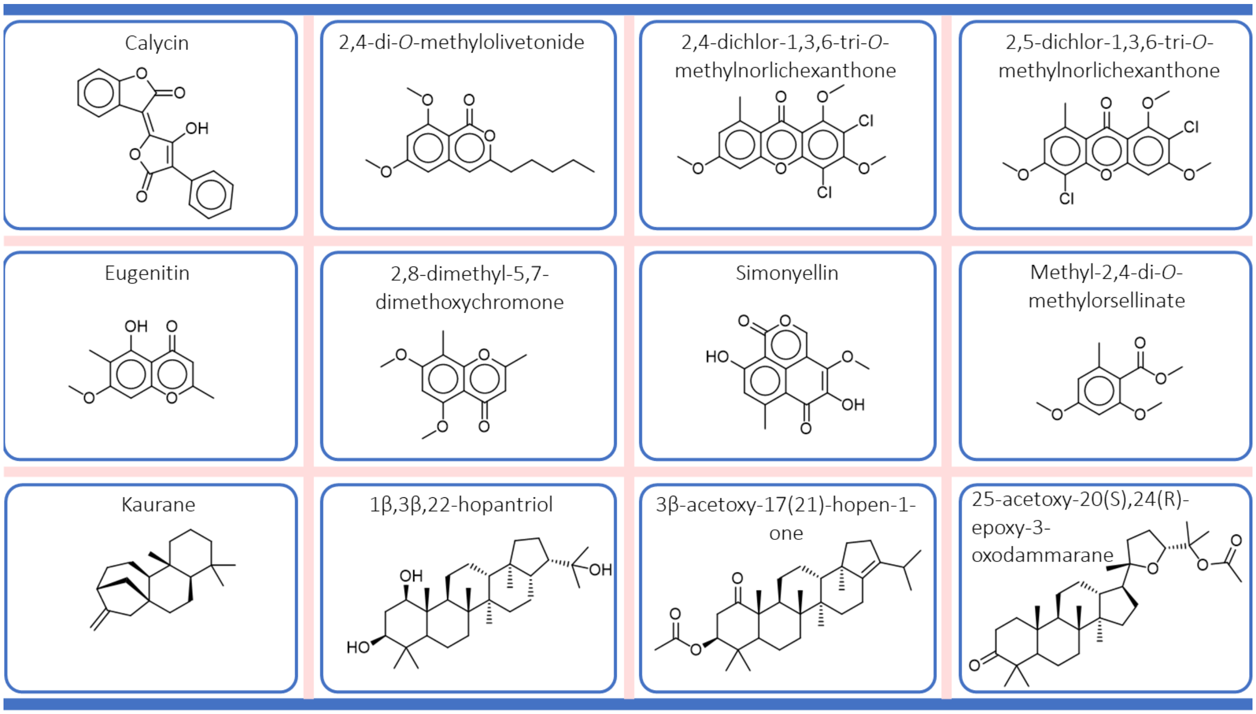


**Calycin**: No neutral node was formed but three singleton ions were dereplicated with the LDB: an [M-H]⁻ at RT = 199 s, another [M-H]⁻ with RT = 167.88, and an [2M-2H+Na]⁻ at RT = 245 s. Given the difference in retention time, it could not be expected that these adducts form a neutral (rt error = 10 s). These ions probably belong to calycin cis- and trans-isomers that are produced spontaneously by pulvinic acid derivatives.

**2,4-di-*O*-methylolivetonide**: only one ion was found belonging to this molecule (a $[2M+Na]^+$). Triangulation was impossible and no neutral was formed.

**2,4-dichlor-1,3,6-tri-*O*-methylnorlichexanthone**: no ions from the LDB for this molecule were found in the processed data.

**2,5-dichlor-1,3,6-tri-*O*-methylnorlichexanthone**: no ions from the LDB for this molecule were found in the processed data.

**Eugenitin**: no ions from the LDB for this molecule were found in the processed data.

**2,8-dimethyl-5,7-dimethoxychromone**: a single ion was found for this molecule ($[2M+Na]^+$), triangulation was impossible, no neutral was formed.

**Simonyellin**: a single ion was found (potentially $[M+H]^+$, low quality, not dereplicated). No neutral was formed.

**Methyl-2,4-di-*O*-methylorsellinate**: a single ion present in the LDB ($[M+H]^+$ at RT = 172 s), found by *MolNotator* to be an $[M+H+CH_3OH]^+$ of a molecule. The molecule also displays an $[M+H]^+$ adduct with a high cosine score (1.0) between both ions. This could however be a false positive: the first adduct might very well be an $[M+H]^+$ instead of an $[M+H+CH_3OH]^+$, and the second one, an $[M+H-CH_3OH]^+$ instead of an $[M+H+]^+$. Differentiating adducts like $CH_3OH$ from their neutral loss in the source is challenging as they are virtually the same. This type of false positives can be expected when there are few adducts for a molecule and the adducts present might be confused with the equivalent neutral losses (**Figure S8A**).

**Kaurane**: one single ion found ($[M+H]^+$), no neutral could be formed.

**1$\beta$,3$\beta$,22-hopantriol**: no ions from the LDB for this molecule were found in the processed data.

**3$\beta$-acetoxy-17(21)-hopen-1-one**: its sole ion from the LDB was found ($[M+NH_4]^+$, RT = 338 s), triangulated as a $[M+H+CH_3CN]^+$ by *MolNotator* in a relatively large molecular cluster, although with low cosines. The *MolNotator* interpretation seems the most likely here, and the LDB entry is probably an error (**Figure S8B**).

**25-acetoxy-20(S),24(R)-epoxy-3-oxodammarane**: the single ion present in the LDB was found, no other adducts were found, no neutral was formed.

Figure S8: Ion from the LDB that were annotated differently by *MolNotator*. A: case of false positive for methyl-2,4-di-*O*-methylorsellinate. B: case of a potential error in the LDB for 1$\beta$,3$\beta$,22-hopantriol

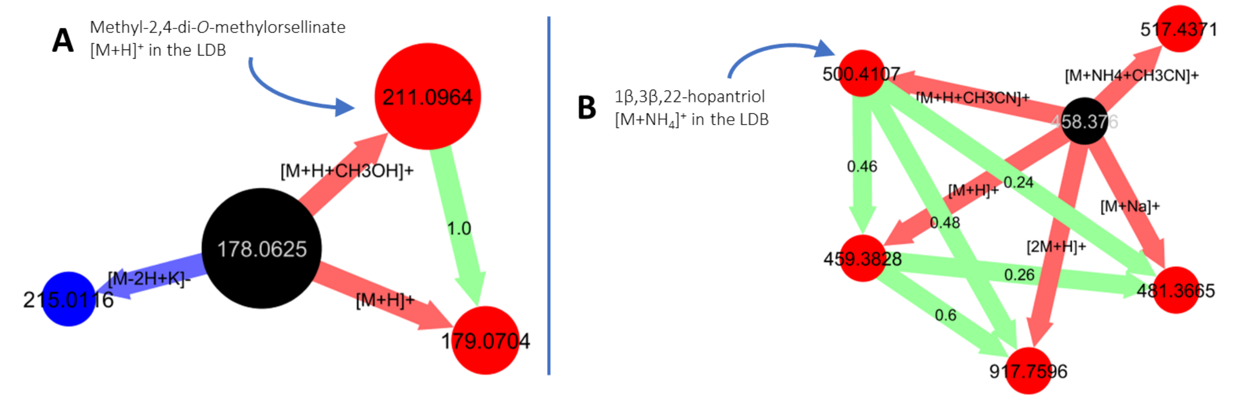
